# Whole-body plethysmography revisited

**DOI:** 10.1101/2021.02.24.432470

**Authors:** Swen Hülsmann, Amara Khan, Liya Hagos, Martin Hindermann, Torsten Nägel, Christian Dullin

## Abstract

Whole-body plethysmography (WBP) is an established method to determine physiological parameters and pathophysiological alteration of breathing in animals and animal models of a variety of diseases, reaching from pulmonary diseases to complex neurological syndromes. Although frequently used, there is ongoing debate about what exactly is measured by whole-body-plethysmography and how reliable the data derived from this method are? Here, we designed a simple device that can serve as an artificial lung model that enables a thorough evaluation of different predictions about and around whole-body plethysmography. Using our lung model, we confirmed that during WBP two components contribute to the pressure changes detected in the chamber: 1) the increase of the pressure due to heating and moistening of the air, termed as conditioning, during inspiration; 2) changes of chamber pressure that depend on airway resistance. Both components overlap and contribute to the temporal pressure-profile measured in the chamber or across the wall of the chamber. Our data showed that a precise measurement of the breathing volume appears to be hindered by at least two factors: 1) the unknown relative contribution of each of these components; 2) not only the air in the inspired volume is conditioned during inspiration, but also air within the residual volume and death space that is recruited during inspiration. Moreover, our data suggest that the expiratory negative pressure peak that is used to determine the so called “enhanced pause” (Penh) parameter is not a measure for airway resistance as such but rather a consequence of the animal’s response to the airway resistance, using active expiration to overcome the resistance by a higher thoracic pressure.

## Introduction

Unrestrained whole-body plethysmography (UWBP) is a widely used physiological technique to analyze breathing behavior in mice. The technique is not only used for the analysis of lung and airway function as such (DeLorme & Moss, 2002; Vondráková *et al*., 2006; Lim *et al*., 2014; Dullin *et al*., 2016) but it is also very helpful for the analysis of physiological or pathological changes of central respiratory control (Gaultier & Gallego, 2008; Chao *et al*., 2010; Abdala *et al*., 2014; Wegener *et al*., 2014; Hulsmann *et al*., 2016; Janc *et al*., 2016).

In principle, two modes of WBP can be discerned, pressure plethysmograph (PWBP) and flow plethysmograph (FWBP) (Lomask, 2006). In 1955, James E. Drorbaugh and Wallace O. Fenn devised a sealed chamber to measure tidal volume in infants (Drorbaugh & Fenn, 1955). They used a “barometric method” (PWBP) utilizing the pressure changes resulting from the warming of the inspired air and cooling during expiration (Drorbaugh & Fenn, 1955) and assumed that at the end of inspiration the inspired air has been warmed to body temperature and humidified to saturation. Based on this assumption the tidal volume is calculated. In contrast, recording chambers devised for FWBP not only are open to the environment and are “perfused” by a bias flow of air or other gas mixtures (Chand *et al*., 1993), but utilize the principle of a pneumotach to measure the flow from inside to the outside of the chamber -or vice versa- and the tidal volume is calculated via integration of the flow (Lomask, 2006). Despite this successful story of UWBP, there is a long standing and still ongoing debate of whether it allows defining of the tidal volume in small animals correctly (Epstein & Epstein, 1978; Epstein *et al*., 1980; Enhorning *et al*., 1998; Lundblad *et al*., 2002; Zhang *et al*., 2014). Moreover, FWBP is often used to quantify the effect of bronchoconstrictors on airway resistance using the enhanced pause (Penh) parameter, which is also constantly debated (Bates *et al*., 2004; Lomask, 2006; Pichavant *et al*., 2007).

In recent study, we have compared the measurement of tidal volume (V_T_) from anaesthetized mice using flow plethysmograph (FWBP) with 3D reconstructions of X-Ray micro-computer tomography (µCT) data, and found that V_T_ calculated from FWBP are smaller as compared to volumes derived from the 3D reconstructions of µCT (Khan *et al*., 2021). Currently, it is unclear why µCT and FWBP data do not match. This difference might be due to technical limitations of the µCT data, such as the difficulty to delineate airspace from micro-vasculature and tissue due to limited resolution of the images (Khan *et al*., 2021). On the other hand, it was also speculated that the uncertainty in FWBP to define the zero-flow point limits the accuracy of the FWBP (Khan *et al*., 2021).

Considering the ongoing discussion about the validity of lung volume-measurements by WBP, we devised a simple mechanical model of a lung, which allowed to perform a series of experiments comparing the tidal volumes derived from FWBP with the known volume of the model, to assess the use of Penh in the model, and compared the model to *in vivo* FWBP and µCT data.

## Methods

### Ethical approval

For in-vivo lung function measurements, male C57Bl/6 mice were used. Mice were kept under 12/12 light/dark cycle and had access to water and food ad libitum. All animal procedures were performed in compliance with the guidelines of the European Directive (2010/63/EU) and the German ethical laws and were approved by the administration of Lower Saxony, Germany. Permission was granted by the Nieders. Landesamt für Verbraucherschutz und Lebensmittelsicherheit (LAVES, approval number G15.1747).

#### X-ray based lung function

ray based lung function (XLF) measurements were performed with a small-animal *in vivo* µCT (QuantumFX, PerkinElmer) allowing acquisition of a 20 x 20 mm^2^ field of view (Dullin *et al*., 2016; Khan *et al*., 2021). XLF was recorded at speed of 30 s^-1^ using an X-ray tube voltage of 90 kV and a tube current of 100 µA. The radiographs were sequenced and exported as TIF-series. Synchronization between µCT and FWBP was performed as described (Khan *et al*., 2021). The transparency of the lung was measured by using the “Plot Z-axis Profile” function of ImageJ software. A region of interest (ROI) was set to the level of the diaphragm in expiration. Data was exported as ASCII format and imported to LabChart 8.0 (Adinstruments). For removal of low frequency intensity changes resulting from the X-ray tube, a “smoothed” intensity trace (median filter over 101 samples) was subtracted from the raw trace. Processed image data and the UWBP data was exported IGOR Pro 6.37 (Wavemetrics) for resampling (10 kHz) and reimported into LabChart for final offline analysis.

### Design of a lung model

For our lung model, we aimed to model as many aspects of the lung as possible (Figure 1). For the airspace of the lung cavity we used a rubber bulb (neoLab®, Heidelberg), as a temperature source – resembling the circulation – we placed the rubber bulb on top of a heating foil (*thermo* Flächenheizungs GmbH, Rohrbach). Expiration was managed by squeezing the elastic rubber bulb with the lever of a servo motor (SG90, TowerPro, Taiwan). The servo motor was controlled by the analog output of the Powerlab 8/35 using a single-board microcontroller (Arduino, Torino, Italy). Inspiration was achieved by lifting the lever from the bulb and expiration by squeezing the bulb again. Different airway resistances were modelled by attaching a modified luer lock cannula tips (B. Braun, Melsungen) to the outlet of the rubber bulb (Figure 1B).

**Figure 1:**
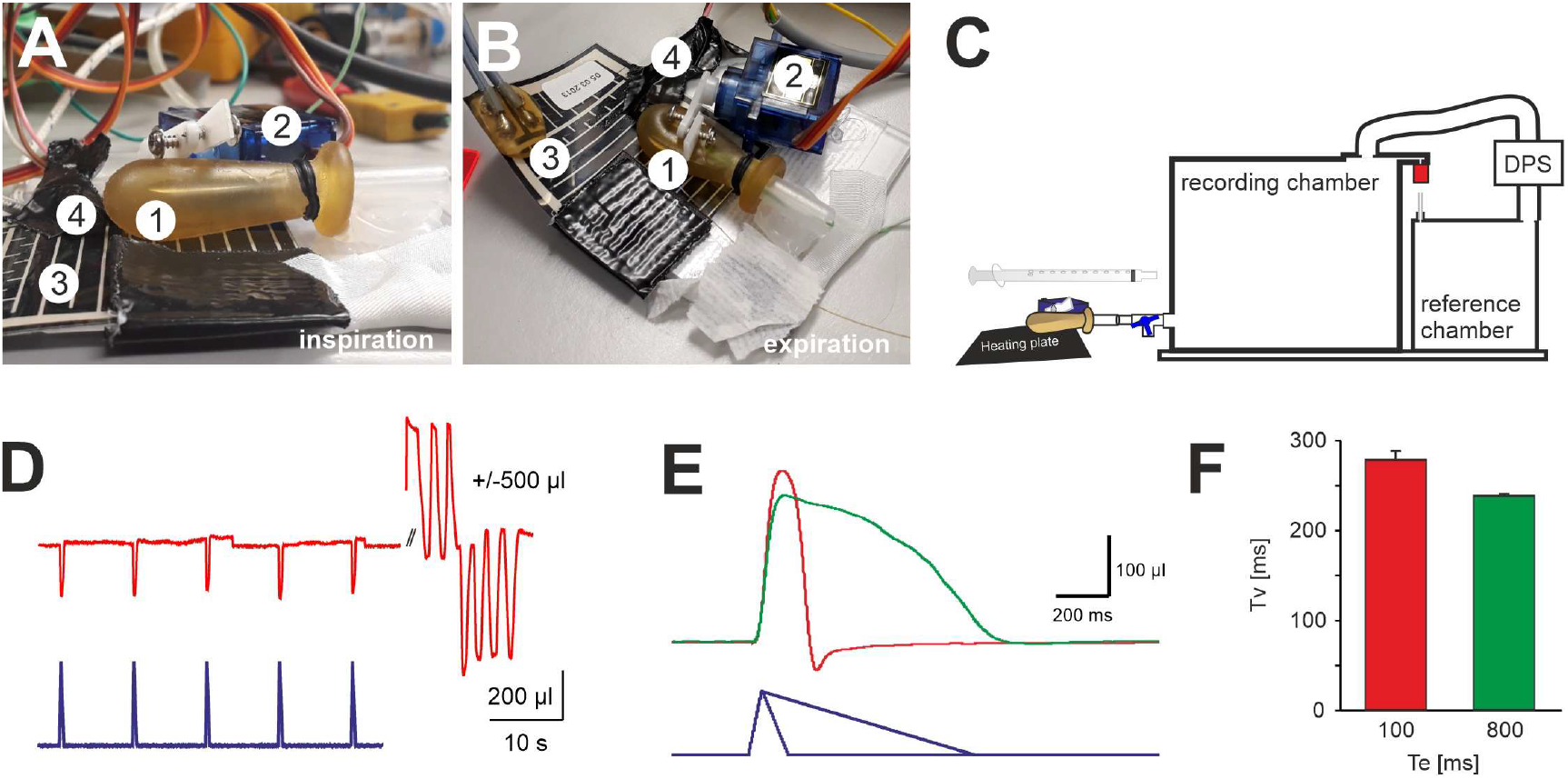
Measurement of the tidal volume (V_T_) that is exchanged during a “respiratory” cycle of the artificial lung model (ALM). **AB:** Design of the ALM. **A:** lateral view of the ALM in inspiration (1) latex bulb, (2) servo motor with lever, (3) heating plate, (4) temperature sensor. **B:** top view of the ALM. **C:** schematic drawing of the experiment. To test the volume the ALM, it was connected to the plethysmography chamber. **D:** recording of 5 cycles. Each inspiration (T_I_ 50ms, T_E_ 100ms) leads to a reduction of the pressure in the chamber (red trace). Blue trace analog out control signal. The pressure changes were calibrated by injection and withdrawal of 500 µl of air using a 1 ml syringe attached to the same port. **E:** Enlarged (inverted) view of the calibrated pressure change (volume) from a respiratory cycle of the ALM with T_I_ 50 ms, T_E_ 100 ms (red) and T_I_ 50ms, T_E_ 800ms (green). **F:** Average data of the volume that is exchanged during one respiratory cycle. DPS = differential pressure sensor.

### Whole-body plethysmography chambers

We used two versions of plethysmography chambers. (i) For the µCT, a chamber was designed that fitted into the gantry of the µCT system (Khan *et al*., 2021). In brief, we recorded the pressure differences between a cylindric recording chamber (220 ml) and a reference chamber (50 ml) by a differential pressure sensor (DPS, Board Mount Pressure Sensor, 0-1” H_2_O, 20 mV, 16 VDC supply; Mfr. No: INCH-D-MV, Amphenol Cooperation Wallingford, CT, USA). (ii) For the test with the artificial lung model (ALM), we used a UWBP-chamber for mice (Hirrlinger *et al*., 2019). Here, the pressure difference between the recording chamber (1180 ml) and a reference chamber (130 ml) was captured by a differential pressure transducer (Validyne Engineering, Northridge, CA, USA). The raw signal passed through a sine wave carrier demodulator (CD-15; Validyne Engineering). Digitization (1 kHz sampling rate) was performed with an analog-digital interface (PowerLab 8/35; ADInstruments) and LabChart-software (ADInstruments). When the chamber was used in the flow whole-body plethysmography (FWBP) mode, a negative bias flow of 150 mL min^-1^ was introduced using a CO_2_/O_2_ sensor (ADInstruments). The low pass filter properties of the chamber were determined by the time constant of the pressure decay, which could be altered by attaching silicon tubing (inner diameter 1.5 mm, outer diameter 3.5mm) of different length (0.5 mm to 300 mm) to the air inlet of the chamber (see below). The time constant was defined by briefly obstructing the air-inlet and then measuring the decay of the pressure signal after releasing the obstruction (LabChart Peak Analysis Module). All cables were fitted through the lid of the chamber, which was sealed by polymer clay (BORT therapeutic putty standard, BORT GmbH, Weinstadt).

### Calibration of the chambers

In the FWBP-mode the chamber represents a pneumotach that can be calibrated via the bias flow (Hamelmann *et al*., 1997). An independent calibration was repeated for each time constant using a two-point calibration (LabChart). The raw flow signal was band pass filtered off-line (0.5 – 20 Hz) to reduce noise, and then integrated for calculation of the tidal volume. We used the standard integral settings of the “Integral Channel Calculation module” of the LabChart-software (use of all data points, reset each cycle whereby the integral is reset each time the source signal passes through zero to a positive value).

Additionally, the chamber was used in the closed mode to perform pressure whole-body plethysmography (PWBP). Therefore, the inlet and the outlet of the chamber were closed by a luer lock plug or a three-way valve, respectively. The latter allows to reset the baseline, which shifts during the experiment due to the constant heat that is added by the ALM. Here, 500 µl air were injected and/or withdrawn by an insulin syringe to calibrate the pressure changes.

### Video analysis of the ALM

To exactly define the timing of the changes of the parts of ALM with respect to the pressure measured in the chamber we recorded slow-motion videos (240 frames per second) from the ALM using a smartphone (iPhone SE, Apple, Cupertino, CA, USA). For synchronization of the video with the chamber-pressure signal, the light of LED (FiberOptoMeter, NPI electronics, Tamm, Germany) was controlled by the analog-output, which also controlled the servo motor, and the coupled into a 200 µm Ø optical fiber, which was taped to the chamber outside within the field of view of the smartphone (Figure 4). The video was exported to an imaging software (*Fiji* distribution of ImageJ; (Schindelin *et al*., 2012)). ROI were placed over the LED-fiber tip, the lever of the servo motor and the latex bulb of the ALM to measure the intensity (gray scale). The signal from the photomultiplier tube (PMT) and the intensity profile of the fiber/LED allows exact matching of the signals from the video and the A/D interface.

**Figure 4:**
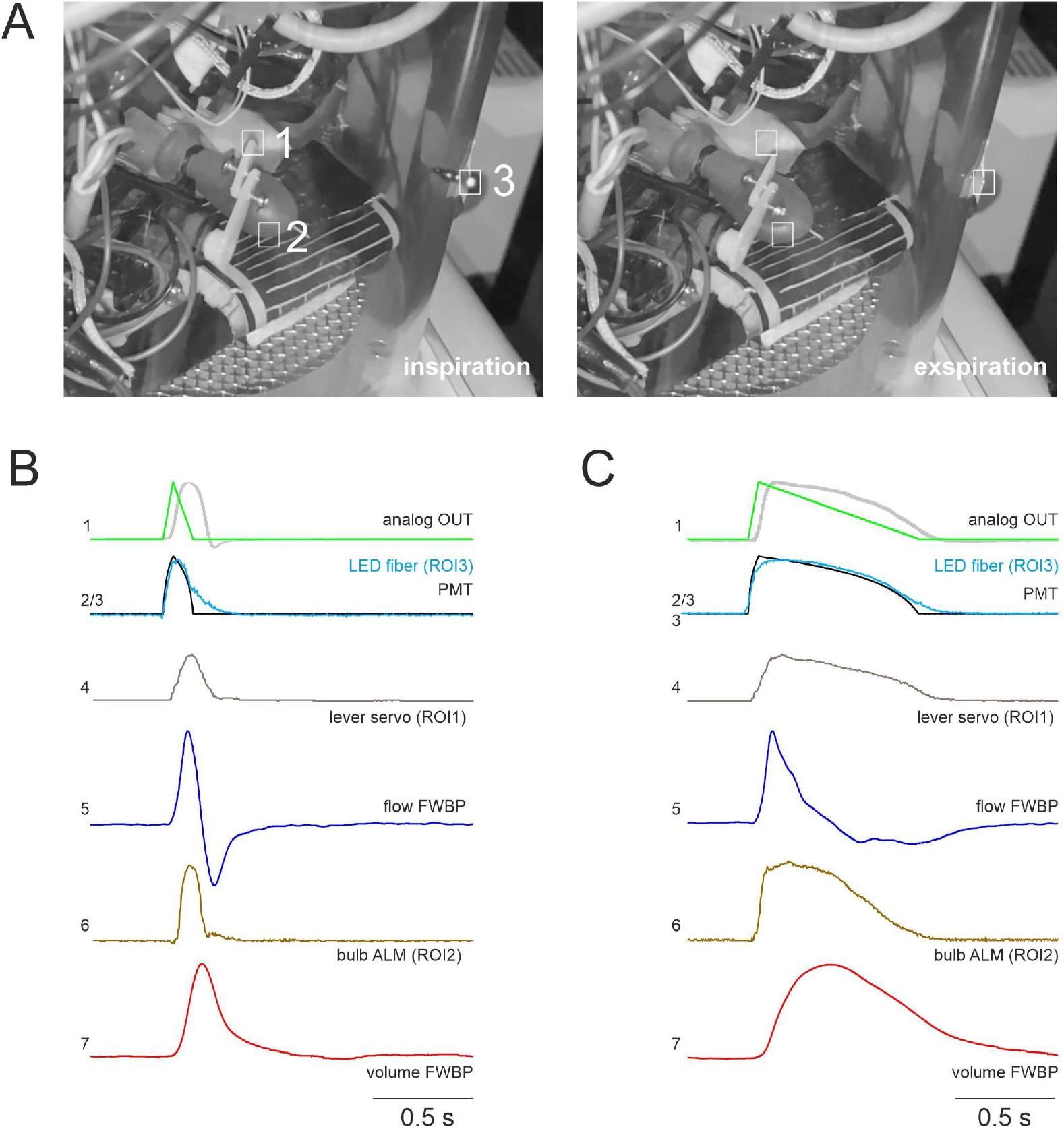
Flow whole-body plethysmography (FWBP) of the artificial lung model (ALM). **A**: Two frames; inspiration (left) and expiration (right) from a video of the ALM during the FWBP taken at 240Hz. Region of interests (ROI) were placed on the servo motor lever of the ALM (ROI1), on the bulb of the ALM (ROI2) and on the tip a light guide that allowed the synchronization with the plethysmography measurement (ROI3). **B:** Timing of different parameters of the ALM with 50 ms duration of the inspiration (T_I_) and 100 ms expiration (T_E_). (1) Green trace analog out from the digitizer. The gray trace represents the inverted pressure measured in the bulb according to the measurement shown in Figure 1. (2) PMT signal of the LED (3) that was driven from the analog out and was also the source for the light signal that was recorded by the video. Graphical overlay of the to signals allows exact matching of the signals from the video and the A/D interface (4) Intensity of the ROI 1 from the servo motor lever indicating the response of the ALM. (5) Flow signal of the FWBP. (6) Intensity of the reflex from the bulb of the ALM. (7) Volume of the FWBP. **C:** Timing of the FWBP-parameters (see B) from ALM recording with a T_I_ of 50 ms and a T_E_ of 800 ms. Traces are averages of at least 3 consecutive cycles.

### Temperature and humidity adjustment of volume

For the experiments with the mechanical lung model temperature or humidity was performed inside the µCT plethysmography chamber using a Bluetooth Hygrometer/Thermometer (WS07; Shenzhen Seven Like Co., Ltd). Additionally, temperature could be measured at the surface of the heating foil or inside rubber bulb with a wire probe (GTF 300) attached to a digital thermometer (GTH1170; Greisinger, Regenstauf, Germany) or to the Powerlab A/D interface (ADinstruments). Additionally, we used a FLIR A655sc infrared camera (FLIR Systems, Inc., Wilsonville, OR 97070, USA) to determine the temperature gradients in our ALM (supplemental figure). To correct for the changes of the pressure that relate to the temperature-dependent expansion and saturation of the air in the lung or ALM, a calculation of the volume adjustment factor was performed.

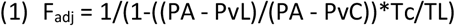

with P_A_ = atmospheric pressure [mmHg], Pv_L_ vapor pressure of water in lung [mmHg]; Pv_C_ vapor pressure of water in chamber [mmHg]; T_L_ = temperature in lung [K]; T_C_ = temperature in chamber [K] (Drorbaugh & Fenn, 1955; Pappenheimer, 1977; Stephenson & Gucciardi, 2002). T_L_ was measured with a temperature probe placed on the heating foil.

### Data analysis and interpretation

Experiments were repeated up to three times using the model as technical replicates. Since the timing was always the same in the different sets of experiments, traces from one set of experiments could be overlaid to others to compare timing of e.g. flow, pressure and volume. The peak detection algorithm of LabChart was used to determine amplitude, decay and frequency of the plethysmography signal. Results were exported to Excel (Microsoft Corp., Redmond, WA, USA) for further offline calculations and to SigmaPlot 14.0 (Systat Software, Inc; Richmond, CA, USA) for statistical interpretation. To facilitate the data transfer and generation of figures in CorelDraw (Corel Corporation, Ottawa, ON, Canada) averaged data was exported from LabChart (ADInstruments) to IGOR Pro 6.37 (WaveMetrics Inc., Lake Oswego, OR, USA). To isolate the conditioning effect in experiments with different airway resistance, subtraction of traces was performed in IGOR Pro.

## Results

### Tidal volume of the artificial lung model

In a first set of experiments, we defined the tidal volume of our artificial lung model (ALM) in two experimental settings and for two different timings of inspiration to expiration ratio (Figure 1). The pressure changes induced by a single respiratory cycle of ALM were compared to known volumes that were injected by a syringe attached to the same port of the chamber. In the first setting, the inspiration time (T_I_) was set to 50 ms and the time of expiration (T_E_) was set to 100 ms, while in the second settings, resembling lower expiratory efforts, T_E_ was set to 800 ms. With the first setting (T_E_ = 100ms) a T_V_ of 286.0 ± 5.5 µl and second setting (T_E_ = 800ms) a T_V_ of 239.5 ± 5.7 µl was measured (n=3; p<0.001; Figure 1EF).

### Conditioning of the inspired air produces significant pressure changes

The warming and saturation with vapor during inspiration is expected to increase the pressure in the connected lung-chamber system during inspiration (Drorbaugh & Fenn, 1955). Accordingly, during expiration the drop of pressure in the chamber is a result of cooling of the expired air. However, it had been debated, whether this conditioning of the air in the lung contributes to the pressure changes that are recorded in WBP (Enhorning *et al*., 1998). Indeed, if the airway resistance is high, one would expect that the chamber pressure rises at the beginning of inspiration without conditioning. Since any resistance slows the airflow into the expending lung, there will be a pressure gradient with a negative pressure in the lung that is accompanied by an increase in the chamber pressure (Enhorning *et al*., 1998).

Therefore, we tested the contribution of both components in our lung model (Figure 2). Using the FWBP mode of the chamber, we recorded the chamber pressure in two conditions. In condition 1, the rubber bulb of the ALM was heated, in condition 2 the heating plate was turned off. When there was no cannula attached to the ALM, a substantial increase of the chamber pressure was measured only when the heating was turned on but not when it was off (Figure 2D). This observation is in line with the concept that warming of the air inside the ALM during inspiration induces an increase in the pressure inside the chamber.

**Figure 2:**
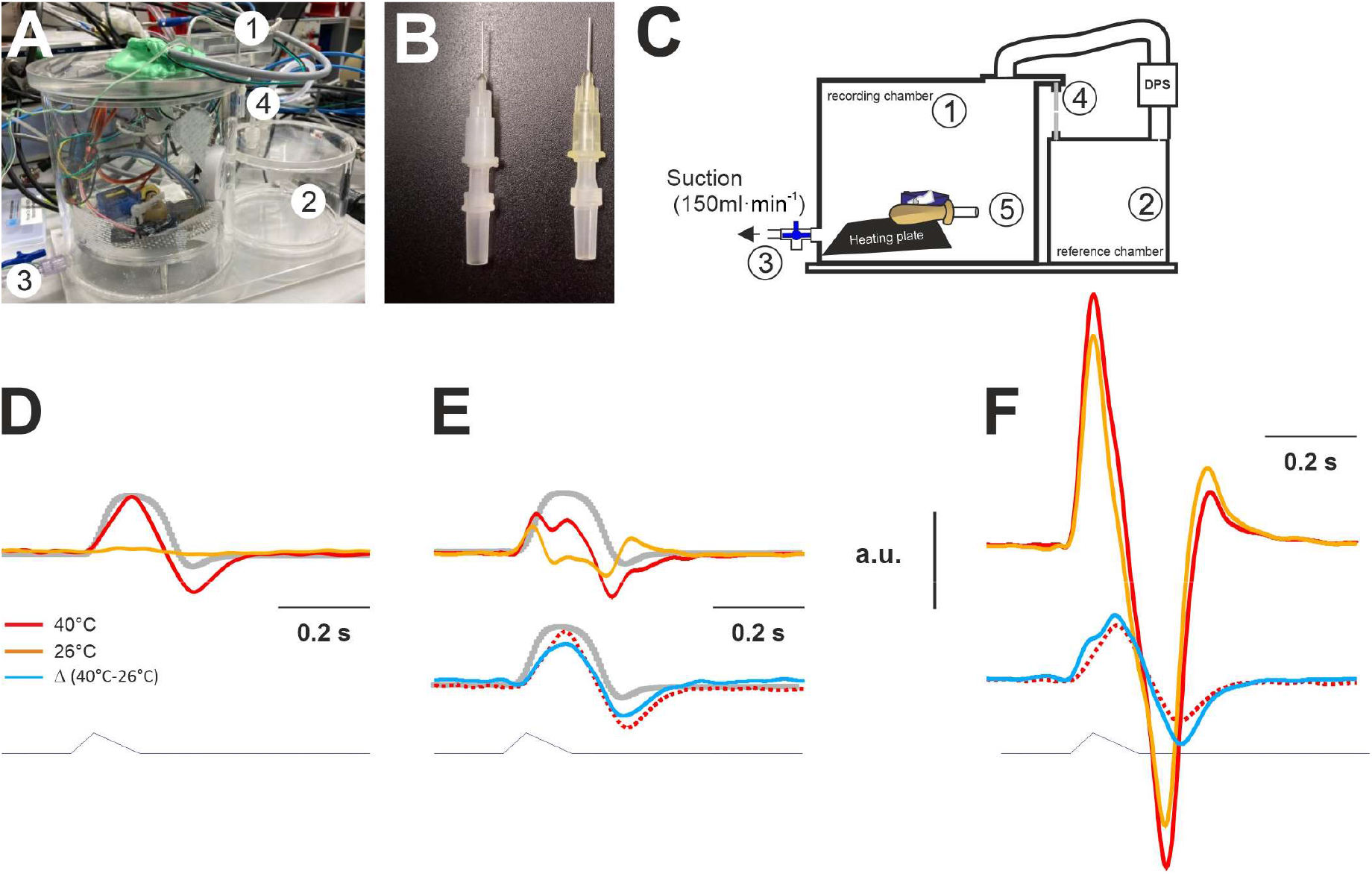
Test in ALM in the plethysmography chamber. **A:** Photograph of the lung dummy in the FWBP plethysmography chamber. Cables passing through the lid of the chamber, secured and sealed by polymer clay (1). **B:** Different cut luer-lock cannulas were used to adjust the resistance of the ALM. **C:** Schematic drawing of chamber design. The flow through mode was used by applying constant suction of 150ml/min to the outlet port (3) with a 3-way-valve. At the inlet port (4), the flow was limited by a defined resistance, a silicon tube (τ 50 ms). Pressure changes were recorded with a differential pressure sensor (DPS) between the recoding chamber and a reference chamber (2). **D-F:** Analysis of the effect of the different degrees of airway resistance. To mimic low airway resistance no cannula was attached to “tracheal” opening of the dummy lung (5). To apply additional resistance either a cut 20G (yellow, **E**) or a cut 27G (white, **F**) canular was attached to the luer adapter. **D:** Chamber pressure (non-calibrated) in the no-resistance situation. The lower trace represents the analog control signal for the micro servo motor (T_I_ 50 ms, T_E_ 100 ms). Traces in red are from recordings with the heating plate turned on (40 °C; see Figure 1) and the yellow trace with the heating plate off (26 °C). The chamber pressure follows the lever of the servo motor with a short delay (see also Figure 4). Note that with no temperature added to the ALM, almost no pressure change is measured (yellow trace). **E**,**F:** Effects of addition of airway resistance. **E:** With the first resistance (20 G) the pressure in the chamber changes already in the unheated dummy lung (26 °C), with an obvious difference between the signal at 40 °C (red) and 26 °C (yellow). Subtraction (blue trace) of the 26 °C signal from the 40°C signal reveals the pressure change that is based on warming of the air during the breath. Interestingly, it almost equals the change when no resistance was used (dotted trace). **F:** Using a larger resistance (27 G), the pressure changes are increased, moreover the difference between the signal at 40 °C and 26 °C is not easy to appreciate anymore. However, subtraction (blue trace) of the 26 °C signal from the 40 °C signal reveals that the flow that is based on warming is still very similar. The gray trace (D) resembles the pressure curve shown in figure 1E.

When we placed a 20G cannula at the opening of the ALM, the pressure signal was different. The pressure in the chamber incresed during the initial phase of the inspiration, even in the unheated ALM (26 °C; Figure 2E). The increase of the pressure started earlier and quickly decreased again, followed by a negative peak during expiration (when the bulb was squeezed again; Figure 2E). When the heat was turned on, a two-peak positive pressure change was detected during inspiration and a second negative dual pressure peak during expiration. Subtraction of the two signals resulted in a pressure fluctuation that resembled the pressure fluctuation of the 40 °C lung dummy with no resistance. With an even higher resistance (27 G cannula), the difference between the signal at 40°C and 26 °C appears to be minimal, however, subtraction of the 26°C signal from the 40 °C signal reveals that the component based on conditioning is still comparable.

These data clearly support the concept that there are two overlaying signals in every plethysmography trace: (i) the “conditioning” signal resulting from warming of the inspired air in the lung (no humidification in ALM) and (i) a resistance dependent change of the chamber pressure following expansion of the lung (the rubber bulb) during the initial phase of the inspiration, when the air is obstructed from entering lung from the chamber due to the airway resistance.

### Testing the precision of volume measurement in FWBP

The previous set of data indicated that conditioning of the inspired air alone is sufficient to induce an increase of the pressure in the chamber, which produces a corresponding airflow across the pneumotach in the chamber wall. Next, we tried to determine whether the integration of the flow signal provides a correct volume information. The integration of the flow resulting from cycles with T_I_ 50 ms and T_E_ 100 ms yield a V_T_ of 59.2 ± 4.3 µl (n=3). Adjusted for temperature and humidity this equals an adjusted tidal volume (adjV_T_) of 426.9 ± 30.8 µl (n=3), which is significantly larger as compared to the known volume (286.0 ± 5.5 µl) that is inhaled by the ALM (p=0.018). When increasing T_E_ to 800 ms the difference between the apparent V_T_ (adjV_T_ 819.1 ± 73.3 µl) and the expected V_T_ (239.5 ± 5.7 µl) was even larger (Figure 3).

**Figure 3:**
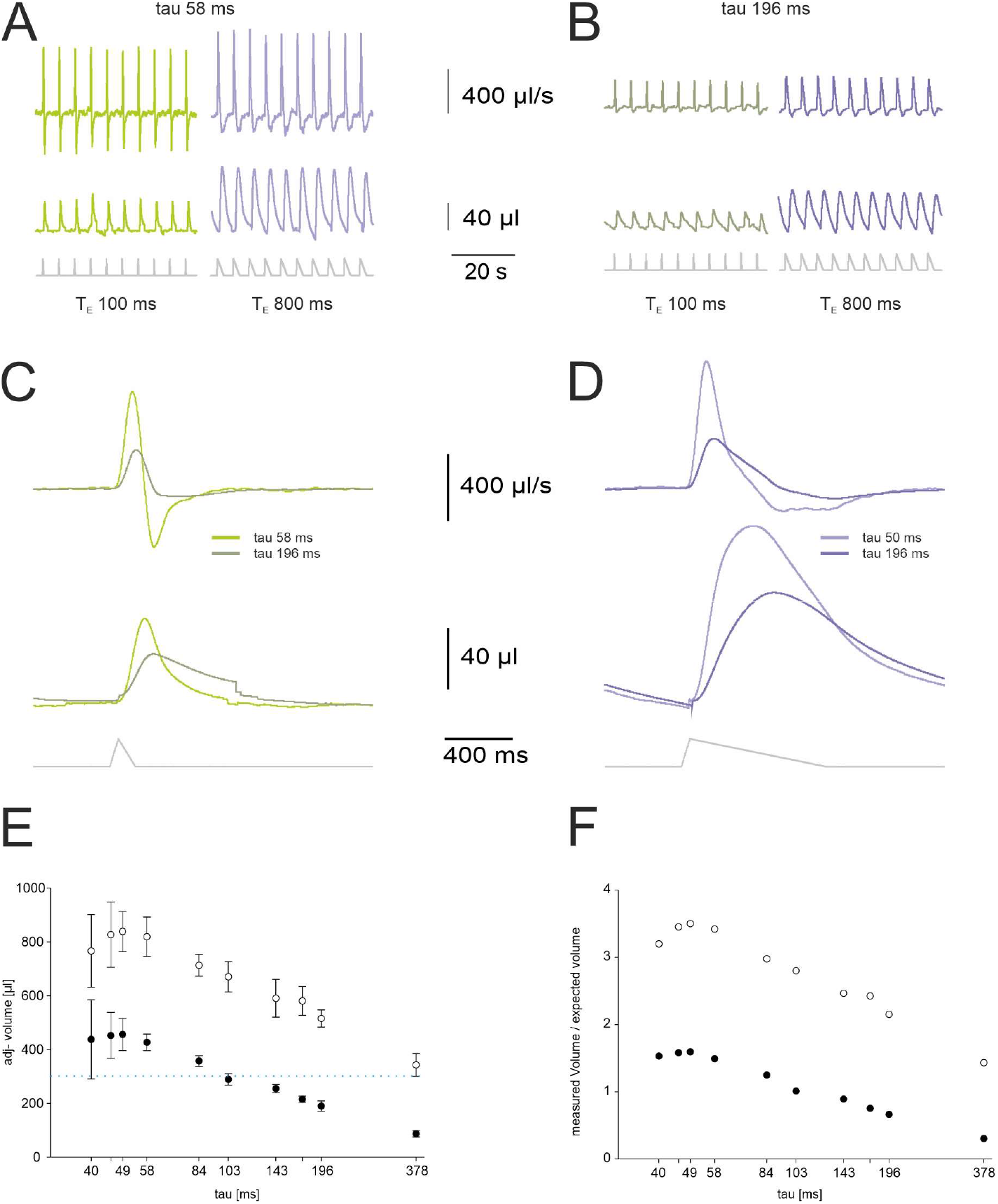
The apparent tidal volume depends on the decay time constant (tau) of the FWBP chamber. **A:** Flow (upper trace) and volume trace from the ALM in a chamber with a pressure decay time constant (tau) of approximately 58 ms for two different durations of expiration (TE 100 and TE T800 ms). **B**: Flow (upper trace) and volume trace from the ALM in a chamber with a pressure decay time constant (tau) of 196 ms for two different durations of expiration (TE 100 and TE T800 ms). **C:** Comparison of the averaged flow (upper traces) and volume traces for respiratory cycles with TE 100 from the ALM in two chambers with different tau. Note that fast time constant leads to a larger apparent tidal volume. With a pressure decay time constant (tau) of approximately 58 ms for two different durations of expiration (TE 100 and TE T800 ms). **D**: Comparison of the averaged flow (upper traces) and volume traces for respiratory cycles with TE 800. The faster tau 58ms leads to a larger apparent tidal volume. **E**: The plot shows the adjusted tidal volume of the ALM in relation to the time constant of the chamber (tau [ms]). Filled circles are from respiratory cycle with TE 100 ms (TI 50 ms), open circles from TE800ms cycles; error bars equal standard deviation. **F**: Comparison of the difference of measured adjTV and known TV of the ALM.

### Influence of the filter properties of the chamber

Next, we tested whether the design of the FWBP chamber, defined by the resistance at the air inlet of the chamber (Table 1; Figure 3), is responsible for the overestimation of adjTV. The volume measurements, described above, were recorded in a flow whole body plethysmography chamber with pressure decay time constant (τ) of 58 ms. However, longer tubes lead to an increase in τ and thus also alters the filter properties of the chamber, since the corner frequencies (fc) of the chamber are defined by τ:

**Table 1:**
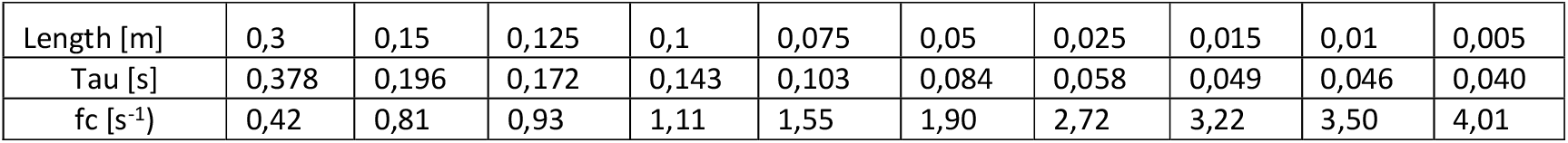
Dependence of low-pass filter corner frequency (fc) from tube length and tau.

|  |  |  |  |  |  |  |  |  |  |  |
| --- | --- | --- | --- | --- | --- | --- | --- | --- | --- | --- |
| Length [m] | 0,3 | 0,15 | 0,125 | 0,1 | 0,075 | 0,05 | 0,025 | 0,015 | 0,01 | 0,005 |
| Tau [s] | 0,378 | 0,196 | 0,172 | 0,143 | 0,103 | 0,084 | 0,058 | 0,049 | 0,046 | 0,040 |
| fc [s<sup>-1</sup>] | 0,42 | 0,81 | 0,93 | 1,11 | 1,55 | 1,90 | 2,72 | 3,22 | 3,50 | 4,01 |

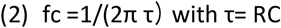

Indeed, when comparing the adjusted tidal volume (adjT_V_) from measurements of the ALM in chambers with different corner frequencies (fc) it was obvious that the fc of the chamber influences the read out of the FWBP. While, measurements with a low fc yield an adjTV that was smaller than the expected T_V_, the T_V_ was overestimated with larger fc chambers. This is, however, what one would expect with the pneumotach in the wall of the chamber: It acts as a low-pass filter (Hirt *et al*., 2008). Although, the interval between two cycles was 2 s (0.5 Hz), the effect of the filter on the amplitude of the pressure signal, however, depends on the rise time T_I_ (50 ms), which is the fastest component of the signal and equal to a frequency component of 10 Hz (the expiration resamples a frequency of 5 Hz (T_E_ 100 ms) or 1,25 Hz (T_E_ 800 ms). Still, even with the highest fc (4.01 s^-1^) used, one expects an underestimation of the adjV_T_ rather than an overestimation. Since, the overestimation of adjV_T_ became larger when using a T_E_ 800 ms as compared T_E_ 100 ms (Figure 3), it appears that the duration of the respiratory cycle (T_E_+T_I_) is responsible for the error (upward shift of curves in Figure 3F).

### Timing of the pressure changes

Next, we tried to determine the exact timing of the changes of the chamber pressure (usually interpreted as flow in FWBP) with respect to the alteration of the ALM, especially the negative pressure in the latex bulb. Interestingly, it became evident that the inspiratory pressure peak (flow signal; blue trace (5) in Figure 4C) in the T_E_ 800 ms situation (heat turned on, no resistance) was very broad and lasted almost 400 ms from start of the pressure change to the end, while it takes about 100 ms to reach the negative pressure peak in the ALM (Figure 1 and 4). This clearly indicates that, while the lung (latex bulb of ALM) is already pushed back to the expiratory position, conditioning of the air in the bulb is continuously happening and thus contributing to the pressure rise in the chamber wall. For T_E_ 100 ms it takes 96.5 ms to build up the pressure, while the chamber-pressure peak still lasts 164 ms (Figure 4B).

### Determination of the tidal volume in the closed chamber configuration

In the next set of experiments, we tested the ALM in the closed chamber without bias flow. Unlike in animals, our ALM does not require any oxygen nor removal of CO_2_. This factor is no limitation regarding the duration of the experiments apply for the ALM. However, this mode is still more prone to pressure artifact and keeping an animal or the ALM inside will lead to a constant increase of the chamber temperature and thus pressure. Although, intermittent opening of the chamber overcomes the pressure rise, the baseline is still difficult to define.

When we measured TV of our ALM in the closed configuration, the calibration was performed by injection of 500 µl air into the chamber. We tested the ALM with and without a 20G cannula as resistance, with T_E_ 100ms or T_E_ 800ms and with and without heat. In the latter situation only a very small increase of the pressure was measured, when no resistance was placed at the opening of the ALM (Figure 5B). However, when the 20 G cannula was used as resistance an increase of pressure was measured during inspiration and a negative pressure peak was measured during expiration when T_E_ was set to 100 ms. Moreover, with a T_E_ of 800 ms the negative pressure peak was missing indicating that resistance not necessarily causes a pressure peak. When the ALM was heated, however, an increase of the pressure and change of the timing of the pressure peaks were measured. Statistical analysis revealed a significantly higher apparent volume (after adjustment for humidity and temperature) for the longer T_E_. Interestingly, the resistance did not change the measured volume significantly (P>0.05; n=4; Figure 5D). Nevertheless, the adjV_T_ was again larger than expected from the initial calibration of the ALM.

**Figure 5:**
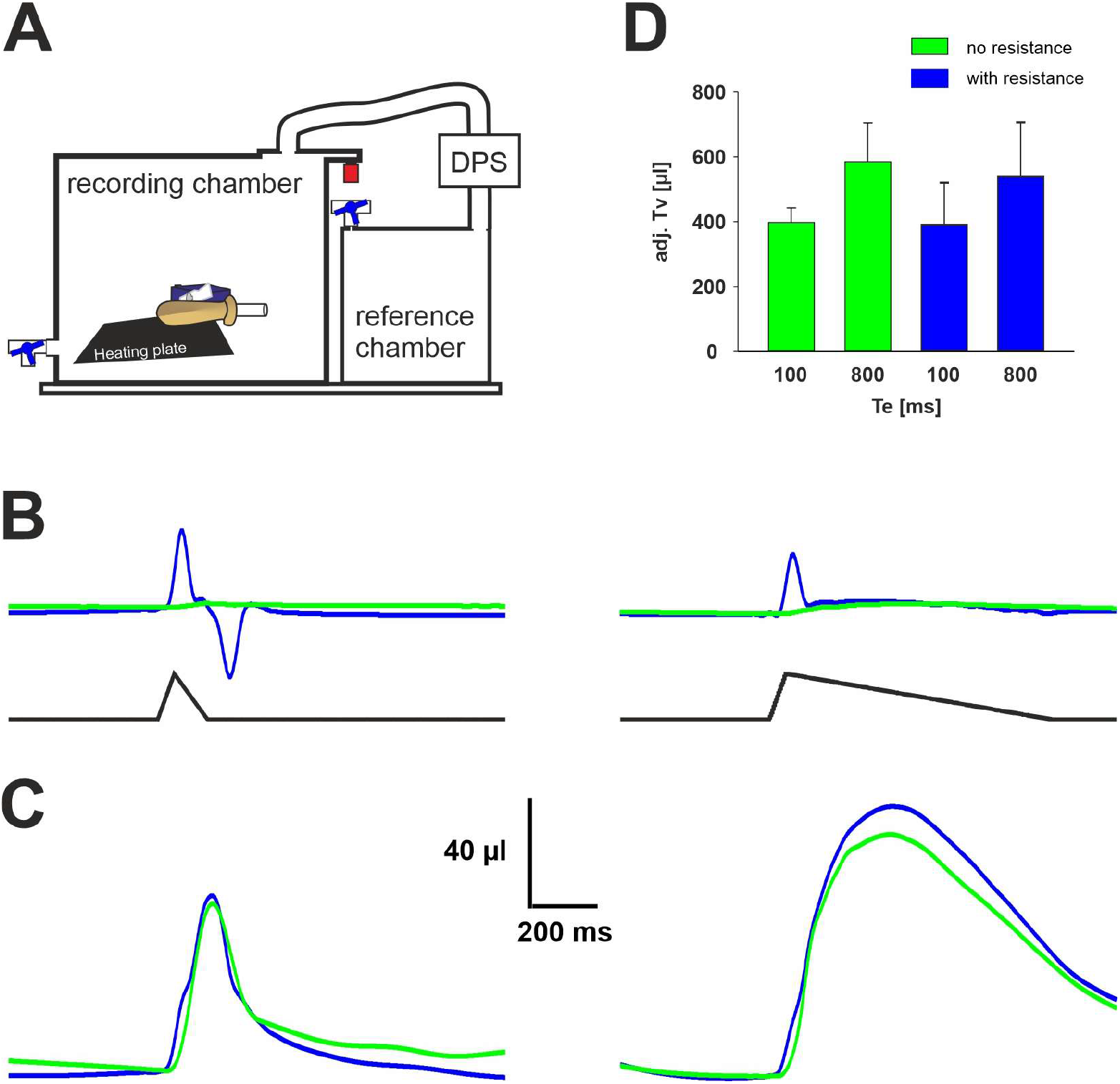
Measurements in the closed chamber (pressure plethysmograph (PWBP) **A:** schematic drawing of the chamber. **B:** raw volume traces from PWBP with the heating plate on (green trace); and with a resistance attached to the ALM (blue trace) for T_E_ 100 ms (right) and T_E_ 800 ms. **C:** raw volume from the same ALM as in (B) with heating turned off. **D:** Bar chart of the basic statistical data of the adjusted volumes. DPS = differential pressure sensor.

### The Penh-parameter

The wave form of the FWBP chamber-pressure changes when the airway resistance is altered (Figure 2). To quantify the waveform-change after e.g. bronchoconstriction the enhanced pause (Penh) parameter was introduced (Hamelmann *et al*., 1997). However, as mentioned earlier the data regarding the validity of Penh is very controversial (Bates *et al*., 2004; Lomask, 2006; Pichavant *et al*., 2007). Our ALM allows to evaluate the Penh concept in a situation, where timing, volume and resistance can be changed under controlled conditions.

In the ALM, increase of the airway resistance is accompanied with an increase of the Penh from 12.64 (no resistance) to 26.51 (27G cannula; Figure 6). When we increase T_E_ to 800 ms, Penh drops dramatical to 0.15 (no resistance) and 0.62 (27G cannula), respectively. This drop is easily explained by the massive reduction of the peak expiratory flow, which in the ALM is a consequence of reduced speed of the servo motor lever. This shows that the PEF is not defined by the resistance as such but rather by the expiratory force of the thorax or in our model of the servo motor. It can be concluded further that the low negative expiratory pressure plateau results from the cooling of the expired air.

**Figure 6:**
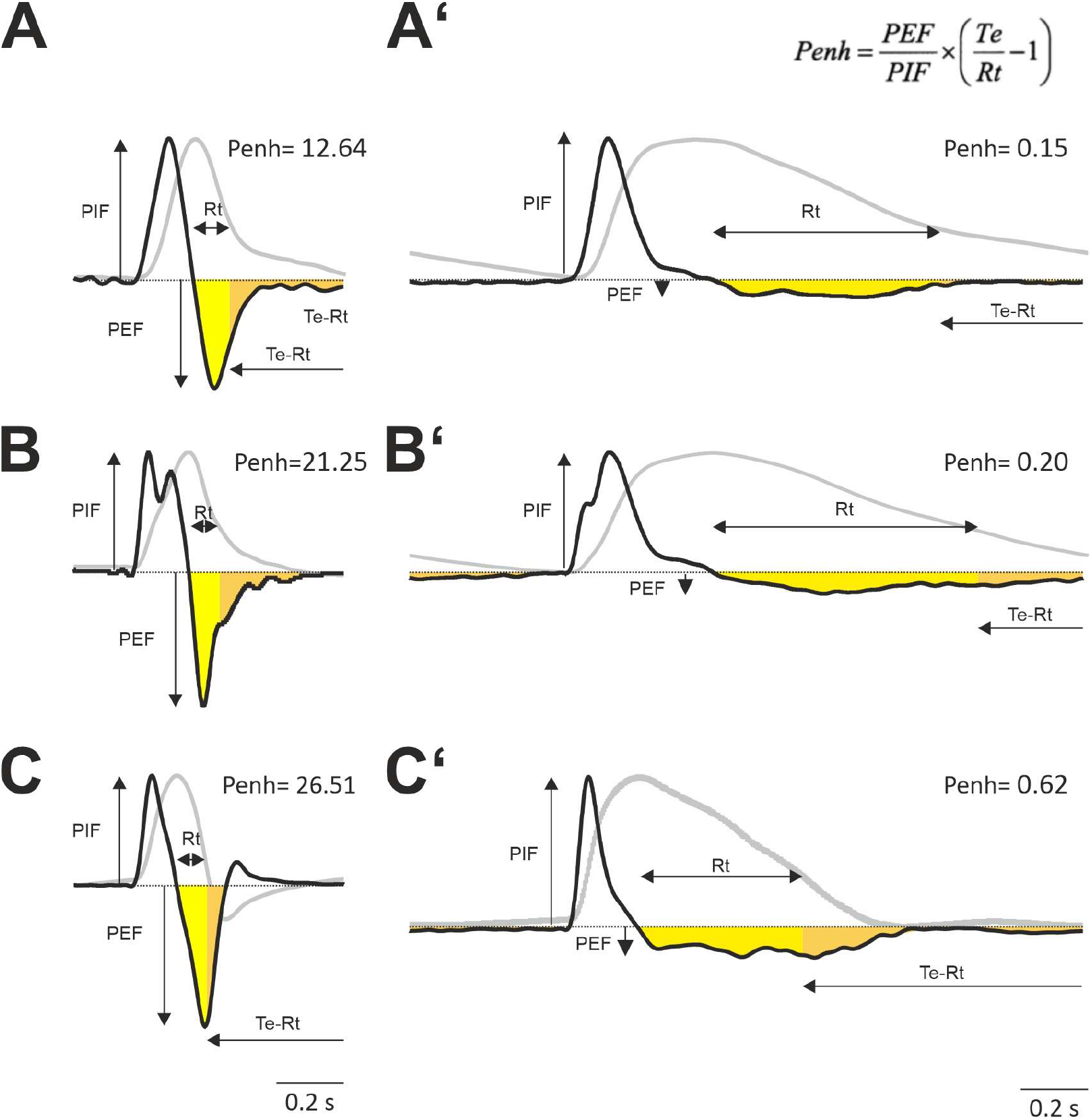
Analysis of the enhanced pause (Penh) paramenter. **A:** Left traces show the flow signal form an ALM (heater on) with no additional resistance (T_i_ 50 ms and T_E_ 100 ms). The gray curve represents the integral of the flow, which equals volume (normalized). A’: same ALM as (A) with T_E_ changed to 800 ms. **B**,**B’:** Measurements as in (A and A’) with a 20 G resistance. **C**,**C’**: Measurements as in (A and A’) with a 27 G resistance. The parameters for Penh are peak inspiratory flow (PIF); peak expiratory flow (PEF); expiratory time (Te), which equals the interval between peak of volume trace and beginning of next inspiration; time to expire 65% of the volume (Rt). In all experiments the respiratory rate was 0.5 s^-1^; traces are average data from 5 to 10 respiratory cycles.

### Effect of anesthesia on airflow and volume changes

The data of our lung model suggest that the flow during expiration depends on the force introduced on the lung rather than on the airway resistance. Indeed, it would be interesting to know, how these results relate to the in vivo settings in mice. Earlier, we have observed a discrepancy between the timing of the lung-volume change during expiration and the pressure in the FWBP recording suggesting that, during isoflurane anesthesia, bronchoconstriction (Eastwood *et al*., 2002; Eilers *et al*., 2010) is resulting in a slower decay of the chamber pressure (Khan *et al*., 2021).

To get additional insight into this phenomenon, we compared the pressure trace from FWBP and the XLF-derived X-ray transmission of the lung in mice with different depths of anesthesia, and thus reflecting different levels of bronchoconstriction (Eastwood *et al*., 2002; Eilers *et al*., 2010). Indeed, when increasing the depth of isoflurane anesthesia, we observed a reduction of respiratory rate and an increase in the amplitude of the inspiratory peak and the opposite effect when isoflurane was washed out. In these experiments it became obvious that during more shallow levels of narcosis indicated by higher respiratory rates (f_R_) the X-ray transmission XLF-curve and volume FWBP-curve are overlapping and that the difference of the duration of the expiration T_E_ is small (Figure 7). However, when depth of anesthesia is increased - as indicated by lower f_R_ - both waveforms become different and the difference between FWBP (fT_E_) and XLF (xT_E_) significantly increases. The T_E_-difference (fT_E_-xT_E_) for shallow levels of narcosis (f_r_ 144.7 ± 24.4 min^-1^; n=5) measured 32 ± 16.9 ms and increased to 200.9 ± 156.8 ms under deep anesthesia (f_r_ 58.4 ± 18.8 min^-1^; n=11; 4 animals; significance p<0.05). These data indicate that isoflurane reduces expiratory pressure drop in FWBP and airflow, while retaining the passive and elastic properties of the thorax and lung, hence any change in the XLF remains rather unaffected.

**Figure 7:**
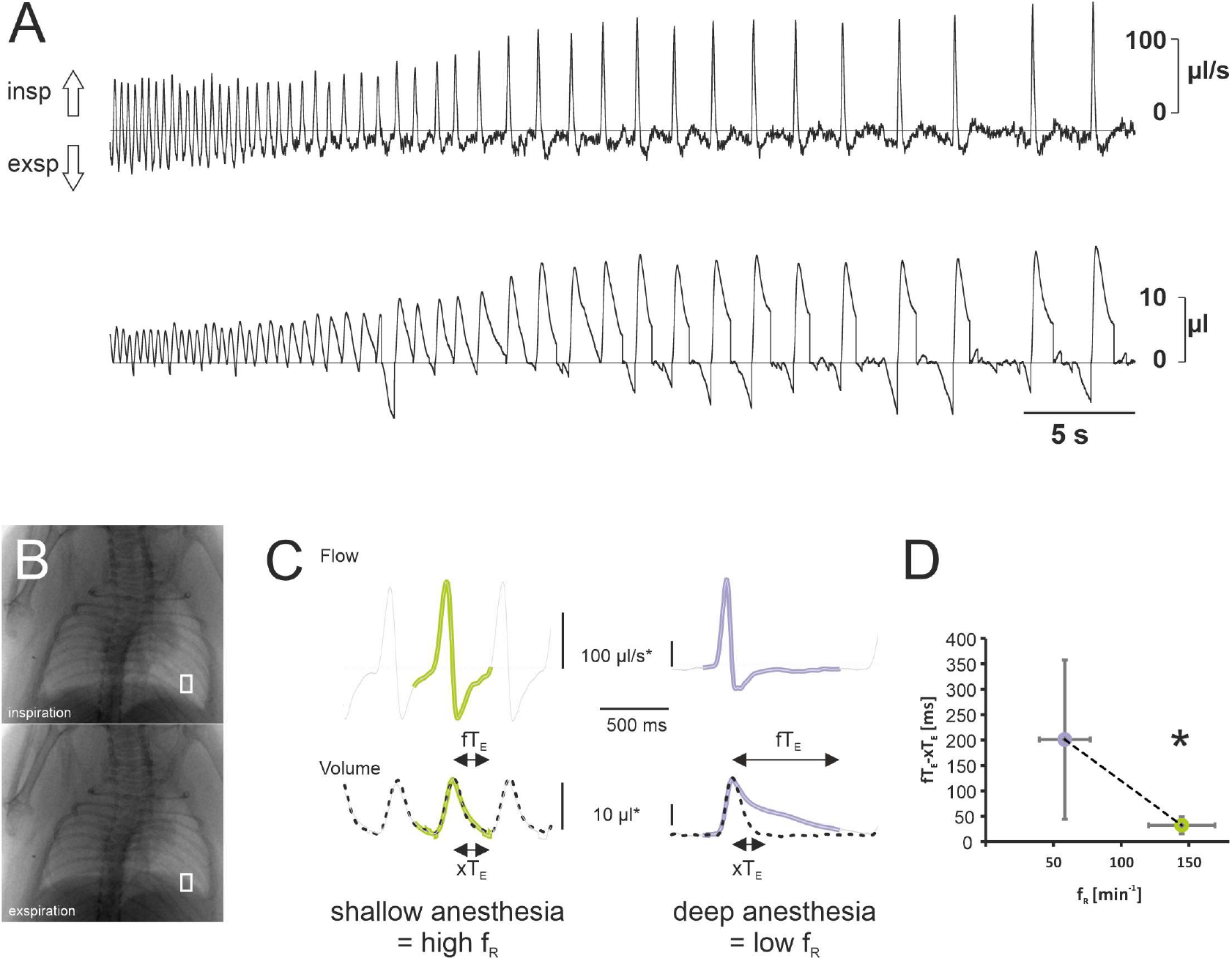
Effect of isoflurane on airflow, tidal volume and XLF-based lung transparence. **A:** Flow whole-body plethysmography (FWBP) trace from an anesthetized mouse. Inspiratory concentration of isoflurane was increased shortly before the recording started. Upper trace shows the flow, while the lower trace shows the integral of the flow over time which represents the volume (both parameters were not corrected for humidity and temperature; see methods). **B:** Example of X-ray-based lung function (XLF) images taken in inspiration and exspiration. **C:** comparison of XLF-based X-ray transmission (dotted line) and FWBP-based flow (upper trace) and volume changes (lower trace) during shallow (green) and deep anesthesia (purple). **D:** Comparison of the arithmetic differences of the duration of expiratory (T_E_) between FWBP (fT_E_) and XLF (xT_E_). Statistical analysis reveals a significant reduction of the difference (fT_E_-xT_E_ [ms]) when the respiratory rate (f_R_) becomes faster (asterisk indicates significance p<0.05).

## Discussion

In principle, our artificial lung model supports the concept that in whole-body plethysmography pressure changes resulting from the warming of the inspired air and cooling during expiration (Drorbaugh & Fenn, 1955) contribute to overall pressure change in the plethysmography chamber and thus to the calculated tidal volume. However, we identified considerable discrepancies between the known tidal volume of the ATM and the adjusted tidal volume (adjT_V_) recorded in both in the FWBP and PWBP configuration. We can exclude that this discrepancy results from the filtering properties of the chamber since it is evident both in the FWBP (Figure 4) as well as in the PWBP configuration (Figure 5). We also can discard a misestimation of the volume adjustment factor (F_adj_; see method) as a substantial cause of the mismatch. F_adj_, required for calculating and correcting the conditioning effect (Drorbaugh & Fenn, 1955; Pappenheimer, 1977; Stephenson & Gucciardi, 2002), is inversely correlated to the body temperature or in the ALM (T_L_) to the temperature of the heating plate. Since the temperature probe for measureing T_L_ was place directly on top of the heating foil, we probably overestimated T_L_ and the actual adjT_v_. vakue should be underestimated. In our model, the air is conditioned as soon as the lever is lifted and inhaled air has contract to the surface of the bulb above heating plate (supplementary figure 1). Since the duration of T_E_ has a positive effect on the overestimation, we hypothesize that constant exchange of air within the bulb of the ALM and from the bulb to the chamber allows conditioning of a larger total volume than what is exactly moved into the lung during inspiration. Assuming that during inspiration all positive pressure changes inside the chamber add up to a flow from inside to the outside of the chamber, the tidal volume read out provided by a FWBP chamber is defined by the integral of all pressure changes. In case the resistance is high, tidal volume will additionally be overestimated by this integration.

However, the opposite effect in the lung might be the case if the contact of the air to the capillary is not long enough to reach vapor saturation and body temperature. This would contribute to the observed discrepancy for the tidal volume between µCT and FWBP (Khan *et al*., 2021).

In FWBP the aforementioned pressure change in the chamber is interpreted as a flow across a pneumotach in the chamber wall. However, it has to be kept in mind that the early inspiratory pressure changes that are seen with high airway resistance, result from the increase of the lung volume, while the airflow into the lung is hindered by the resistance. Likewise, while flow into the lung with high airway resistance is low, the pressure difference (negative in the chamber) is high and vice versa during expiration. Thus, in the condition of high airway resistance, flow across the chamber wall and airway flow are not matched at all.

The latter phenomenon has also important implications on the concept of the enhanced pause (Penh). Penh has been postulated as a measure for the quantification of airway resistance. Penh depends on the quotient of peak expiratory and peak inspiratory (chamber) flow and the time it takes for the peak expiratory volume to drop to 65% (Hamelmann *et al*., 1997). As mentioned above, the validity of Penh, is discussed very controversial, see (Bates *et al*., 2004) and papers therein. Now, our data raises further concerns about the concept behind Penh.

Knowing that the chamber pressure decreases during expiration, a faster and steeper drop of the chamber pressure - as required to increase Penh -is only expected if there is a fast compression of the lung by the lever (in our ALM) or by active expiration (*in vivo*) as seen during vocalization of neonatal mice (Hernandez-Miranda *et al*., 2017; Hulsmann *et al*., 2019). During vocalization of mice, unlike in humans, where the tension of the vocal cords increases airway resistance, calls of neonatal mice depend on the speed of the expiratory airflow (Mahrt *et al*., 2016). Since, the negative chamber-pressure change induced by active expiration during vocalization of mice is accompanied by an increase of flow in the airway, it can be concluded that conditioning, namely the temperature drop during active expiration, is responsible for the pressure drop in the plethysmography chamber during mouse ultrasonic vocalization (Hulsmann *et al*., 2019). In contrast, if the airway resistance is high, without active expiration, a significant pressure drop cannot occur as evident during deep isoflurane anesthesia (Figure 7) or with a T_E_ 800 ms in the ALM. In cases of high airway resistance and active expiration (white cannula in the ALM and T_E_ 100 ms) the airway flow is, however, hindered and the pressure drop is expected to depend mainly on the compression of the lung. And, thus, the volume read out of the pneumotach (integration of the chamber flow) is proportional to the change of the volume in the chamber but not necessary to the volume in the lung. Thus, it is obvious that a fast pressure drop during expiration can occur with and without high airway resistance, and therefore a parameter which strongly depends on the size and timing of this pressure drop, should be used, if at all, only with great care. Further research is required to clarify whether Penh can be used as a measurement for active expiration.

### Limitation of the model

Notably, the lung cavity of our ALM is certainly not mimicking all aspects of the lung. The air ventilated by our model (between 200 and 300 µl; Figure 1) was at the lower range of what we expect in adult mice. Moreover, both total capacity (∼1.7 ml) and residual volume of the model appears to be too large as compared to the mouse. We expect that the single cavity of the bulb as compared to the segmented bronchial tree and alveolar compartment influences the discrepancy between measured and expected tidal volume. In the ALM, the servo motor lever squeezes the bulb such that the two opposing sides of the bulb are put together. Therefore, during expiration, all air is removed from the part that is placed directly on the heater. One can assume that in the animal lung this state is not reached unless during expiration there is an alveolar collapse and atelectasis occurs in pathological situations like ARDS (Schiller et al., 2001).

Further, the tissue elasticity that drives the lungs expiration is missing in our model, and thus, expiration is an active process that is mediated by squeezing of the bulb by the lever of the servo motor. This precludes a simulation of exactly the situation as seen in the mouse where recoil of the lung produces the positive pressure that drives passive expiration. However, the chosen inspiratory time (Ti = 50 ms) allows to constrain the tendency of the bulb to expand.

## Conclusion

In summary, undoubtedly UWBP allows to determine parameters of respiratory rhythm, respiratory rate as well as parameters that define regularity of the respiratory rhythm an also phases of respiratory arrest (Hulsmann *et al*., 2016; Vogelgesang *et al*., 2017; Mesuret *et al*., 2018; Vogelgesang *et al*., 2018). However, for defining tidal volume, UWBP is a weak method. Moreover, our data suggests that the drop of the chamber pressure during expiration, which is used to define Penh, does not only depend on the airway resistance, but also and more importantly on the active force that is driving air out of the lung, which, in the living animal, is defined by active expiration.

## Acknowledgments

The authors thank Anja-Annett Grützner for technical assistance and Johannes Hirrlinger, Carl-Ludwig-Institute for Physiology, University of Leipzig, for discussion and English editing.

## Supplementary Material

**Supplemental figure:**
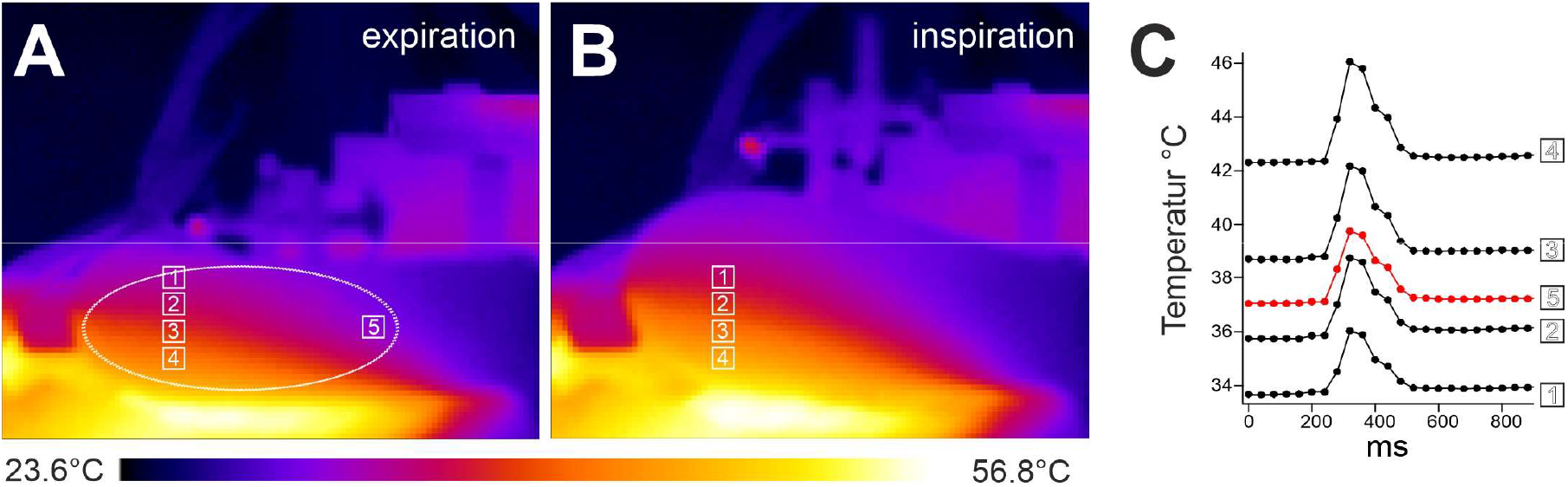
Dummy model of the lung. A-C: Measurement of temperature changes using a thermographic camera (FLIR A655sc) in expiration (A) and inspiration (B). Alterations during a breathing cycle are shown in (C) for 4 individual spots (1-4) and a region of interest (5, red trace) indicating an increase of the bulb temperature during inspiration.

